# Single-shot label-free nanoscopy for quantitative organelle visualization on standard commercial microscopes

**DOI:** 10.1101/2024.12.31.630894

**Authors:** Yugo Inutsuka, Koki Yamamoto, Masafumi Kuroda, Yasushi Okada

## Abstract

Label-free microscopy techniques that overcome the limitations of fluorescence microscopy are increasingly needed in biological research. Quantitative phase imaging has emerged as a powerful label-free method for live cell imaging, particularly at multicellular scales. However, a critical gap exists in achieving easy-to-implement, high-speed, high-resolution microscopy at subcellular scales. Here, we report upgraded polarization-resolved differential phase contrast nanoscopy (UpDPC); this single-shot computational phase microscopy (CPM) enhances resolution and sensitivity to subcellular levels. UpDPC can be readily integrated into commercial fluorescence microscopes, enabling 100 nm gap separation, over 100 Hz imaging via single-shot phase retrieval, and 100 ms computation of phase image during observation. Our results demonstrate that high-performance CPM provides reproducible and comprehensive visualization of various organelles, offering a practical and scalable implementation for basic research and clinical applications.

## Main

Fluorescence microscopy is essential in cell biology; however, it has fundamental limitations in live cell imaging due to phototoxicity, which can damage cells, and photobleaching, which restricts observation time. Additionally, the specificity of fluorescence imaging may prevent comprehensive observation of environmental factors, such as organelle interactions within cells or cellular deformations. Furthermore, the requirement for specific labeling restricts its range of applications. Therefore, it is crucial to explore label-free microscopy techniques that do not rely on fluorescence.

In label-free imaging of cells, phase contrast microscopy, which converts the phase shift of transmitted light into intensity changes, has proven practical for visualizing cells. Zernike phase contrast microscopy (ZPC)^1^ and differential interference contrast microscopy (DIC)^2^ are widely used in cell biology. However, both techniques suffer from limited quantitative phase measurement capability; ZPC is further constrained by reduced spatial resolution and halo artifacts, while DIC exhibits directional dependency in phase detection.

Recently, quantitative phase imaging (QPI), which enables direct measurement of quantitative phase information, has garnered significant attention and has been applied in diverse fields such as biomedicine^3^ and neuroscience^4^. Since QPI images reflect the dry mass distribution of cells^5,6^, this technique has been extensively used at the multicellular scale. Key applications include tracking dry mass increases during the cell cycle^7,8^, estimating cell cycle phases^9,10^, distinguishing normal and abnormal cells quantitatively^11,12^, and quantifying cellular responses to perturbations^13^. Moreover, advancements in machine learning have shown that QPI can be extended to subcellular scales, demonstrating its potential for visualizing organelles similarly to fluorescence microscopy. Previous studies have demonstrated that QPI images can predict the fluorescence images of large phase-shift organelles such as nuclei^14,15^, lipid droplets^15^, mitochondria^15,16^, and lysosomes^16^. Thus, it is anticipated that further improvements in the spatial resolution and sensitivity of QPI would enable the label-free inference of finer organelles or structures.

Among QPI methods, phase retrieval techniques have gained attention. These techniques theoretically formulate the image formation process of microscope optics to compute the phase shifts of samples from captured images. Notable methods include solving the Transport of Intensity Equation (TIE) using defocused images of the sample^17,18^, Fourier ptychography combining images obtained under various illumination angles^19^, and the use of structured illumination patterns in techniques like differential phase contrast microscopy (DPC)^20,21^. These methods, independent of interferometry, demonstrate robustness against external disturbances. However, they typically require multiple image acquisitions under diverse conditions to achieve accurate phase reconstruction (Supplementary Note 1). This necessity compromises temporal resolution and introduces motion artifacts due to calculations using images at different time points. To overcome these limitations, several methods have been developed, enabling phase retrieval from a single acquisition (Supplementary Note 2).

In this study, we developed an upgraded version of a single-shot computational phase microscopy (CPM) that creates illumination patterns using polarization^22^. Through enhancement of both the microscope setup and phase retrieval algorithms, we achieved visualization of delicate intracellular structures. This approach, which can be implemented as a simple extension to a commercial microscope, enables simultaneous observation with conventional fluorescence microscopy. The resolution-optimized setup achieved separation of 100 nm gaps. By leveraging a physical model for phase retrieval, our system provides high-sensitivity capabilities for live-cell imaging and enables real-time phase computation. We demonstrate the system’s capabilities in live-cell imaging applications, establishing its potential as a powerful nanoscopic tool for cell biology.

## Results

### Implementation of Upgraded polarization-resolved differential phase contrast microscopy (UpDPC)

Our UpDPC optical setup (Fig. 1a) requires minimal modifications to a standard bright-field microscope, specifically to the condenser and camera unit. Similar to pDPC^22^, the system requires two additional components: a polarization mask pattern (Fig. 1b) and a polarization-separating camera (Fig. 1c). The UpDPC mask incorporates an opaque line between adjacent polarizing films to prevent light leakage through gaps and simplify the mask fabrication process. A circular block at the optical axis center enhances the detection of spatially high-frequency information at the same light intensity level (Supplementary Note 3). During sample observation, the intensity through each polarization grid of the camera corresponds to images illuminated by different apparent illumination patterns, as shown in Fig. 1d. From these four single-shot images, the phase shifts of the transmitted light through the sample can be calculated (e.g., Fig. 1e).

**Fig 1.**
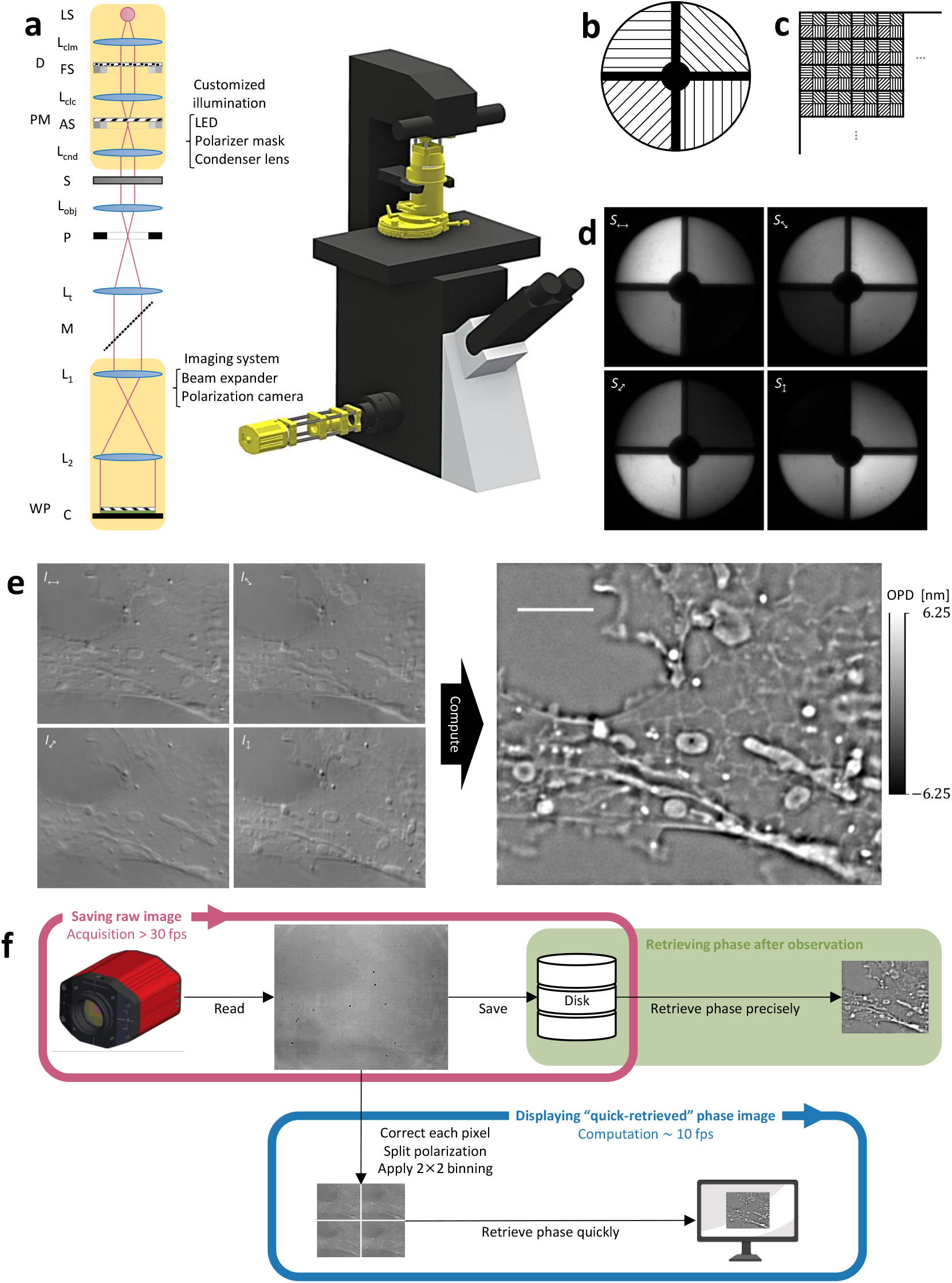
Overview of our microscope. **a**, Setup of our upDPC system on the commercial microscope (left) and 3D scheme of its implementation on the Olympus microscope (right). The additional requirements are highlighted with a yellow color. The light from a light source (LS) is collimated with a collimator lens (L_clm_) and passes through a diffuser (D) and a field stop (FS). A collector lens (L_clc_) collects this light to the illumination aperture plane where the designed polarizer mask (PM) and an aperture stop (AS) are on, and a condenser lens (L_cnd_) collimates this light to illuminate the sample (S). The light passing through the sample is collected by an objective lens (L_obj_) with a pupil (P), expanded by a tube lens (L_t_), and reflected by the mirror (M). Two lenses (L_1_ and L_2_) expand this light, and a camera (C) with a wire-grid polarizer (WP) to resolve a polarization captures this light intensity. **b**, Pattern of polarizer mask inserted on condenser back focal plane. Quadrants of polarizers with four different polarization directions are connected by a cross that blocks light. The center circle also blocks direct light, which primarily carries low-frequency information. **c**, Polarizer array on the polarization camera. Four different direction filter grids are present in each 2×2 pixel. **d**, Apparent illumination distribution for each direction in the polarization camera. In our microscopy, the captured image can be regarded as a mix of images illuminated with these patterns. **e**, Normalized 625 nm LED transmitted light intensity of the COS-7 cells for each polarization direction, obtained with the microscopy (left) and retrieved phase shift image from these transmitted light images (right). The tiled image containing the whole cell bodies is shown in Extended Data Fig. 2. The calibration bar shows the optical path difference (OPD). The scale bar represents the 5 um. **f**, Data acquisition scheme of the code parallelizing the saving thread and display thread. In the saving thread, the image from the camera is read on the global variable named image and saved on the disk as fast as possible. Independently, the image is read, and its phase is retrieved in the display thread. This computation takes about 100 ms per image with a standard PC, which is acceptable when using a microscope. After the observation, a more precise phase contrast with higher resolution with the highest temporal resolution can be obtained.

An additional upgraded feature of this system is its phase retrieval. Accurate phase retrieval demands precise noise handling. Specifically, adding or subtracting independent pixel-wise noise in multiple images increases noise variance. To optimize phase shift estimation despite image noise, we designed an image formation model incorporating not only the optical system but also noise characteristics. Using this model, we perform computations to estimate the most probable phase shift from obtained images, thereby minimizing the noise effect in the phase reconstruction (Methods). Furthermore, accurate modeling of the optical system is crucial. By using a Bertrand lens and observing a glass slide sample (Extended Data Fig. 1c), we precisely determined the spatial distribution of the illumination intensity (Methods, Extended Data Fig. 3). Our model also considers detailed system parameters, such as individual differences in camera pixels and dust on the image sensor.

Furthermore, UpDPC enables real-time phase image generation during microscopy, which is essential for practical applications in high-resolution cell observation. When observing cells using high-resolution CPM, relying solely on the original transmitted light images used for phase retrieval makes it difficult to find focus or detect small phase-shift structures. Real-time phase retrieval during observation is, therefore, essential. However, complex image formation models typically require iterative methods, which can be time-consuming due to the need for convergence. In contrast, our image formation model allows direct computation using only matrix calculations and fast Fourier transforms (FFT) by appropriately setting prior distributions. In practice, after applying 2×2 binning to the polarized images, phase retrieval is completed in 103 ms ± 5.18 ms (mean ± standard deviation of 7 runs, 10 loops each) on a standard PC (Intel Core i7-1360P, 32.0 GB RAM). Using this computation, we implemented a setup that saves camera images to disk at maximum speed while simultaneously displaying the most recent phase images through parallel processing (Fig. 1f). This approach enables not only real-time phase retrieval but also the acquisition of high temporal resolution phase information for post-observation analysis.

The simplicity of the UpDPC setup allows for easy enhancement of spatiotemporal resolution. Specifically, a standard bright-field microscope system can be utilized as is, and commercial condenser lenses, filters, and light sources can be easily integrated. In this study, we enhanced the resolution by using an oil-immersion condenser lens. To achieve a pixel pitch finer than the Nyquist frequency, a beam expander composed of two lenses was placed in front of the polarization camera (Fig. 1a). To improve temporal resolution, it is crucial to implement an illumination intensity that saturates the camera’s dynamic range at its maximum frame rate. The versatility of the UpDPC setup enabled us to achieve this condition by simply replacing the light source with an inexpensive, commercial power LED, achieving sufficient intensity to almost saturate the camera in 8 ms. By narrowing the CMOS camera’s FOV, we achieved 125 Hz imaging, enabling us to capture stepwise vesicle transport driven by molecular motors within cells (Extended Data Fig. 10, Supplementary Video 1).

### Evaluation of UpDPC spatial resolution and live cell imaging

The spatial resolution of UpDPC was evaluated by observing fine structures of known dimensions. We utilized a Siemens star pattern, which is recommended for spatial resolution assessment of coherent imaging systems^23^. Initial evaluation using commercial quantitative phase microscopy targets demonstrated that UpDPC could readily resolve their smallest gap distance of 200 nm (Extended Data Fig. 4).

Given that theoretical considerations suggested the potential for even higher resolution, we fabricated a finer Siemens star pattern on a glass substrate (Methods, Extended Data Fig. 5-6). Using this pattern and applying the concept of the Mutual Transfer Function (MTF), commonly used in Fourier optics (Methods), we demonstrated gap resolution below 100 nm with a 445 nm wavelength illumination (Fig. 2a,b). This achievement aligns with theoretical predictions, as the effective numerical aperture (NA) of UpDPC corresponds to the sum of the NA of the condenser and objective lenses, enabling spatial resolution up to twice the coherent diffraction limit (Extended Data Fig. 7).

**Fig 2.**
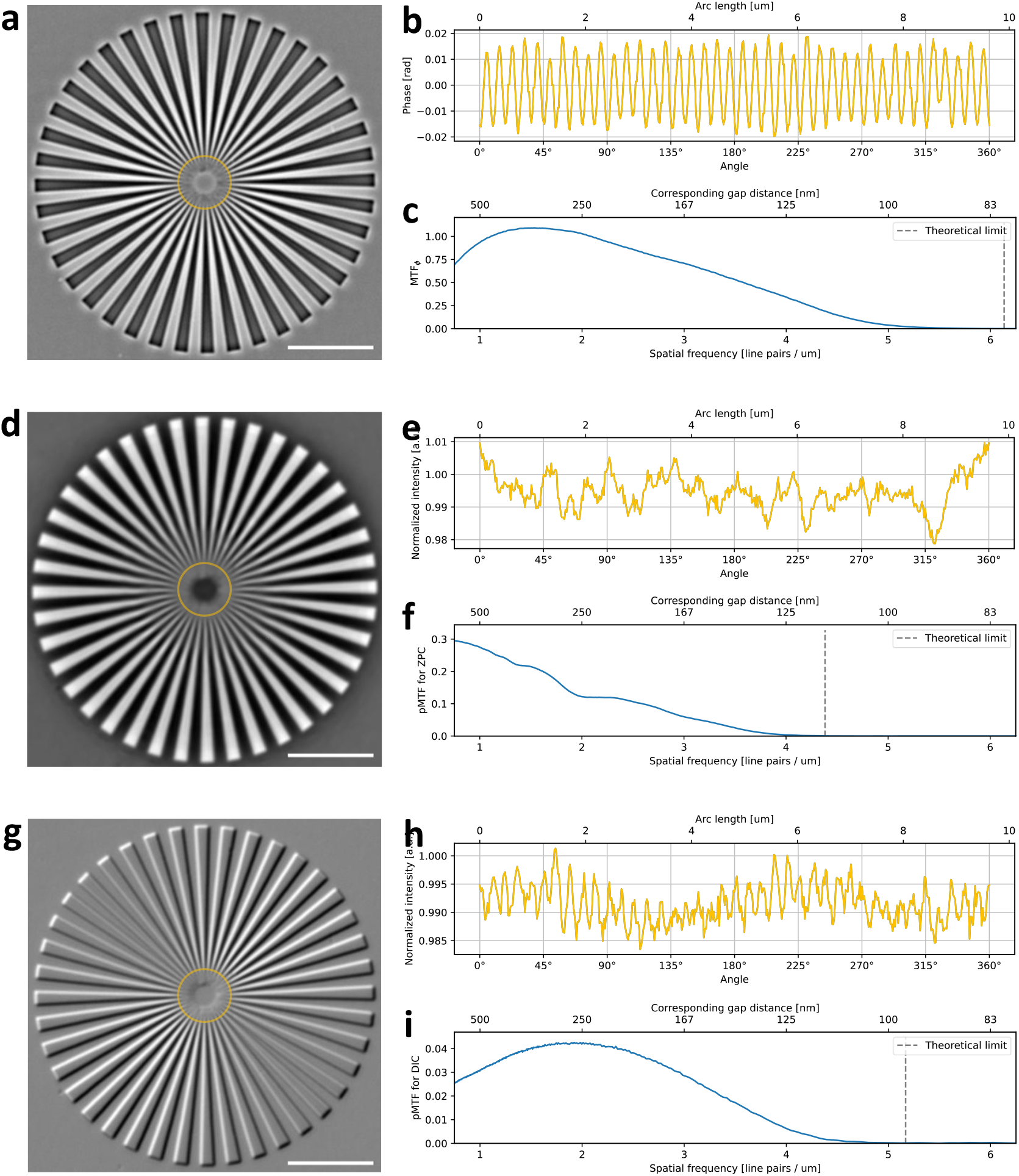
Evaluation of spatial resolution with handmade Siemens-star target. **a**, Retrieved phase image of the handmade Siemens-star target using UpDPC with 445 nm LED. Each adjacent phase spoke on the yellow circle is 120 nm apart. This target is made by carving a fused silica substrate with a refractive index of 1.45 at a carving depth of 50 nm. MilliQ water is filled between this target and the cover glass in observation. **b**, Angular phase shift profile on the yellow circle in **a**. The coordinates are counterclockwise, with the right end at 0°. (The top edge is 90°, the left edge is 180°, and the bottom edge is 270°.) **c**, MTF_*f*_, corresponding to the ratio of the estimated and original phase shifts at each Fourier spatial frequency for **a**. This spatial frequency corresponds to the inverse of twice gaps, i.e., one square wave wavelength. The theoretical resolution limit is overlayed with the dashed line. **d**, ZPC image of the same handmade target as **a**. Intensity is normalized by the background to remove the dust on the image sensor and pixel difference of conversion rate. **e**, Angular intensity profile on the yellow circle in **d**, like **b**. 120 nm gaps are not resolved with the ZPC setup. **f**, The same MTF analysis, substituting the normalized ZPC intensity into the estimated phase shift profile of **c**, with the unit original signal. The MTF curve decays at a lower spatial frequency than UpDPC’s due to the illumination aperture restriction. **g**, DIC image of the same handmade target as **a**. Intensity is normalized by the background similarly to **d. h**, Angular intensity profile on the yellow circle with 120 nm gaps in **g**, like **b**. Phase contrast perpendicular to the prism shear direction (around 135° and 315° in this image) is not transferred to the DIC intensity, causing the orientation dependency of the spatial resolution. **i**, The same MTF analysis substituting the normalized DIC intensity into the phase shift profile of **c**, normalized by the maximum value. Scale bar in **a, d, g** shows 5 um.

We further compared the spatial resolution capabilities by observing the same Siemens star patterns using ZPC (Fig. 2d-f) and DIC (Fig. 2g-i). ZPC requires restricting the illumination aperture, inherently limiting the achievable resolution. Indeed, ZPC was unable to transmit phase shift at smaller spatial frequencies compared to UpDPC. In contrast, DIC converts phase differences into intensity differences along the prism shear direction, making it difficult to detect phase differences in directions nearly perpendicular to the shear. UpDPC, by design, overcomes the limitations of both ZPC and DIC, providing high-resolution phase contrast images without directional dependency.

Furthermore, we evaluated the quantitative accuracy of phase retrieval in UpDPC. Using custom-made Siemens star patterns of different heights, we focused our measurements on a gap width of 300-500 nm, where the transmission efficiency (MTF_*ϕ*_) approaches its maximum (Fig. 2c). We performed measurements with both air and water filling the gaps, thereby varying the refractive index of the medium. The phase shifts measured by UpDPC showed excellent agreement with the theoretical designed values calculated from the pattern heights and refractive indices of the media (Extended Data Fig. 8), confirming the system’s high quantitative accuracy. These results demonstrate that UpDPC provides a reliable tool for quantitative phase measurements at the subcellular level, including vesicles and organelles.

We also compared the performance of these three methods in live-cell imaging (Fig. 3, Extended Data Fig. 9). UpDPC revealed the internal membrane structure of mitochondria by exploiting the refractive index contrast between the matrix and membrane. Additionally, UpDPC clearly visualized density variations within the nucleolus, a structure formed by liquid-liquid phase separation. These results demonstrate that UpDPC enables more intuitive observation of structures that are challenging to discern with ZPC and DIC.

**Fig 3.**
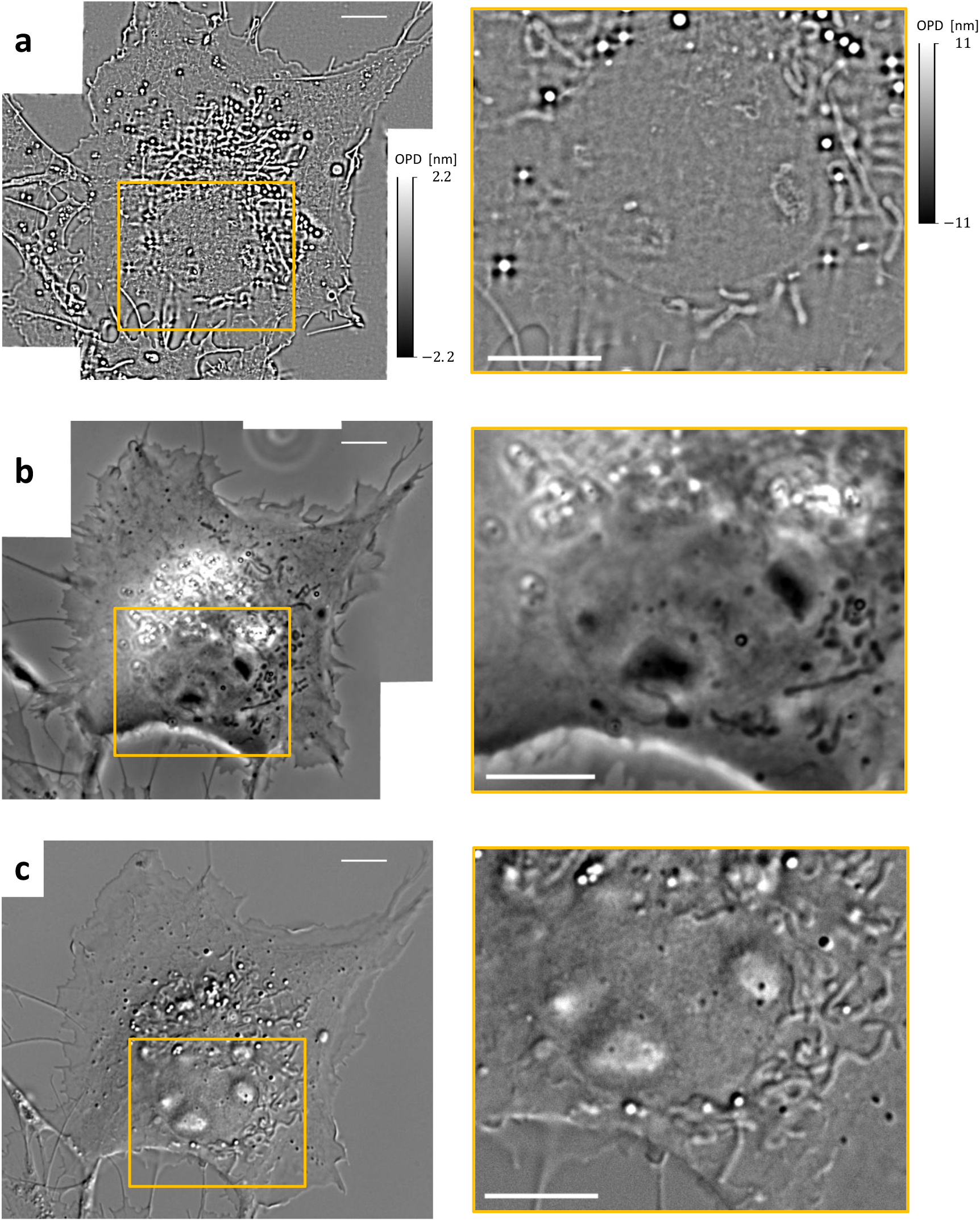
Comparison of the live cell image between UpDPC, ZPC, and DIC. The same COS-7 cell is observed with the UpDPC (**a**), ZPC (**b**), and DIC (**c**) setups using the same 445 nm illumination source and imaging system. The left images are stitched from slightly different time frames to visualize the entire cell. The right images focused the nucleolus on the same FOV as the rectangle on the right images. The images around the nucleus obtained in single exposures are shown on the right. The scale bar shows 5 um.

### Demonstration of UpDPC’s phase shift sensitivity

To validate the high phase contrast sensitivity of UpDPC, we observed microtubules moving on kinesin immobilized on glass in vitro. Microtubules labeled with tetramethylrhodamine were visible under fluorescence microscopy (Fig. 4a). In contrast, single microtubules are challenging to observe with transmitted light microscopy due to the small optical path difference of ∼3.1 nm in water (calculated as (1.64 - 1.33) × 10 nm)^24^ and the small diameter of 25 nm. Nevertheless, UpDPC images of the same FOV successfully reveal these structures (Fig. 4b, left). Additionally, averaging multiple consecutive frames over time reduces the random noise variance proportionally to the number of frames averaged. Thus, we averaged the subsequent 2, 3, or 4 frames captured at 40 ms intervals (with 8 ms exposure time) to enhance the visibility of microtubule phase contrast (Fig. 4b). Moving microtubules were visualized by averaging the UpDPC phase images every 5 frames (0.2 seconds) in Supplementary Video 3 and every 25 frames (1 second) in Fig. 4c. These results demonstrate that fast phase contrast imaging not only enables tracking rapid dynamics but also improves phase contrast sensitivity.

**Fig 4.**
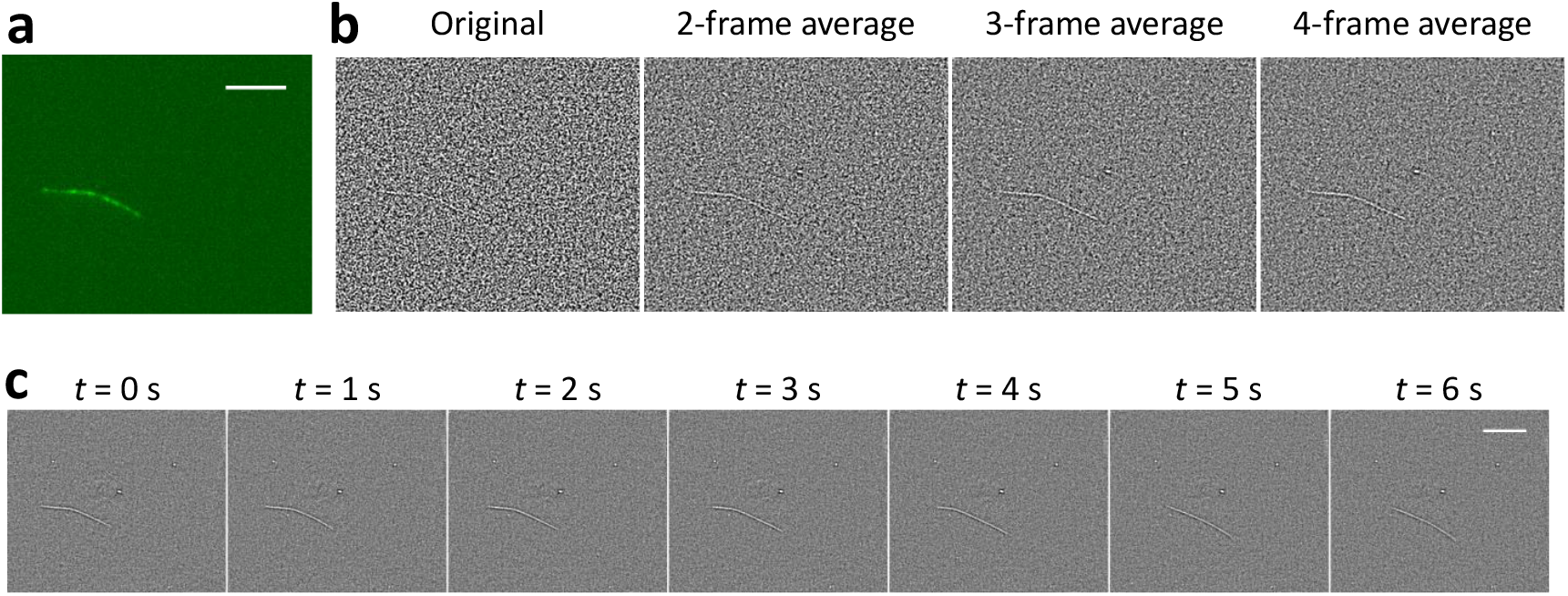
Observation of a single microtubule in gliding assay. **a**, Fluorescence image of a single tetramethylrhodamine-labeled microtubule gliding on kinesin K560 (mouse) attached to the glass surface. **b**, Retrieved phase shift of the single microtubule with 445 nm LED in the same FoV as **a**. From left to right, the images are averages of one, two, three, and four frames. High-speed observation enables the reduction of shot noise by averaging for fast samples. **c**, Time-lapse frames of the retrieved phase of a single microtubule. 25 frames in each 1 second are averaged. Time-lapse movie with averaging 5 frames in each 0.2 seconds is shown in Supplementary Video 5. The phase shift in **b** and **c** shows the contrast between ±0.11 nm optical path difference. The scale bar shows 5 um.

### Observation of organelle dynamics using combined UpDPC and fluorescence imaging

We utilized UpDPC’s capability to acquire fluorescence and phase images in the same FOV, implementing a strategy where fluorescence images are intermittently captured between high-speed UpDPC acquisitions. Structures of interest identified in fluorescence images can then be continuously tracked with UpDPC, with fluorescence imaging reserved for critical confirmation points. This approach minimizes the drawbacks of fluorescence imaging while enabling precise tracking of targeted structures.

Using this fluorescence-phase-fluorescence strategy, we observed the endoplasmic reticulum (ER) in COS-7 cells expressing er-(n2)oxStayGold(c4)^25^, while applying a long-wavelength LED to UpDPC to minimize photodamage. We successfully observed a high-refractive-index structure stretching the ER within the cell (Fig. 5, Supplementary Video 4). This strategy allows high-speed examination of targeted intracellular structures along with their surrounding environment.

**Fig 5.**
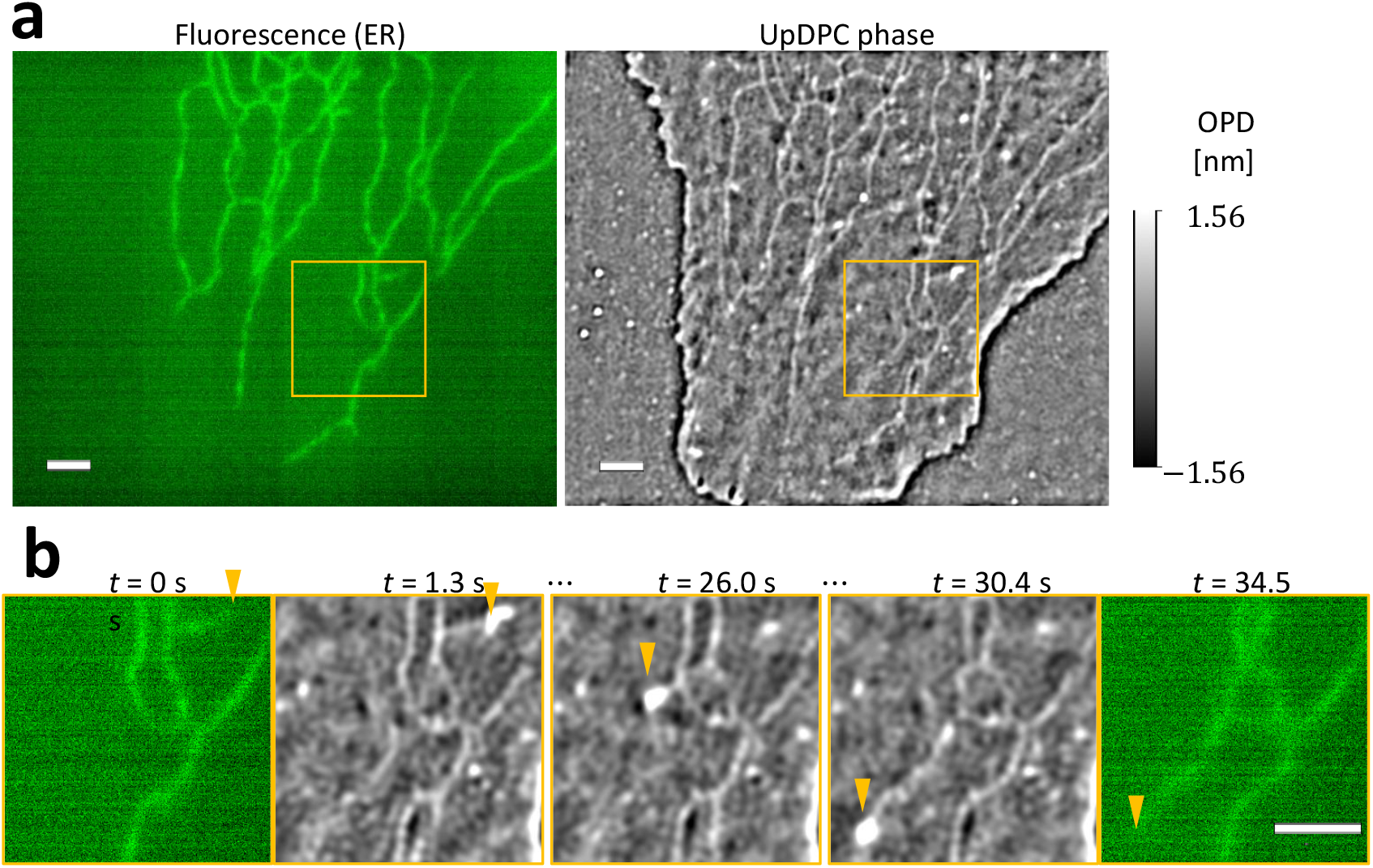
ER dynamics captured by UpDPC system. **a**, Fluorescence image (left) and phase shift image (right) with 625 nm LED of a COS-7 cell expressing er-(n2)oxStayGold(c4). The yellow square indicates the same FoV as the images in **b. b**, Representative frames of the UpDPC phase time series in the same FoV showing extension of ER. The yellow square surrounds the same area in **a**. The yellow arrows mark the extending tip of the ER. The entire track with 30 ms intervals is shown in Supplementary Video 4. The scale bar shows 2 um.

UpDPC’s high resolution and phase contrast sensitivity allow the tracking of various organelles. In addition to previous digital-stainable organelles^14–16^, UpDPC revealed actin bundles, peroxisomes, and stress granules as phase contrast structures (Fig. 6a-c). Although the structure of the Golgi apparatus is challenging to visualize due to the dense surrounding material, we identified its location by detecting regions with a lower refractive index than the surroundings (Extended Data Fig. 11).

**Fig 6.**
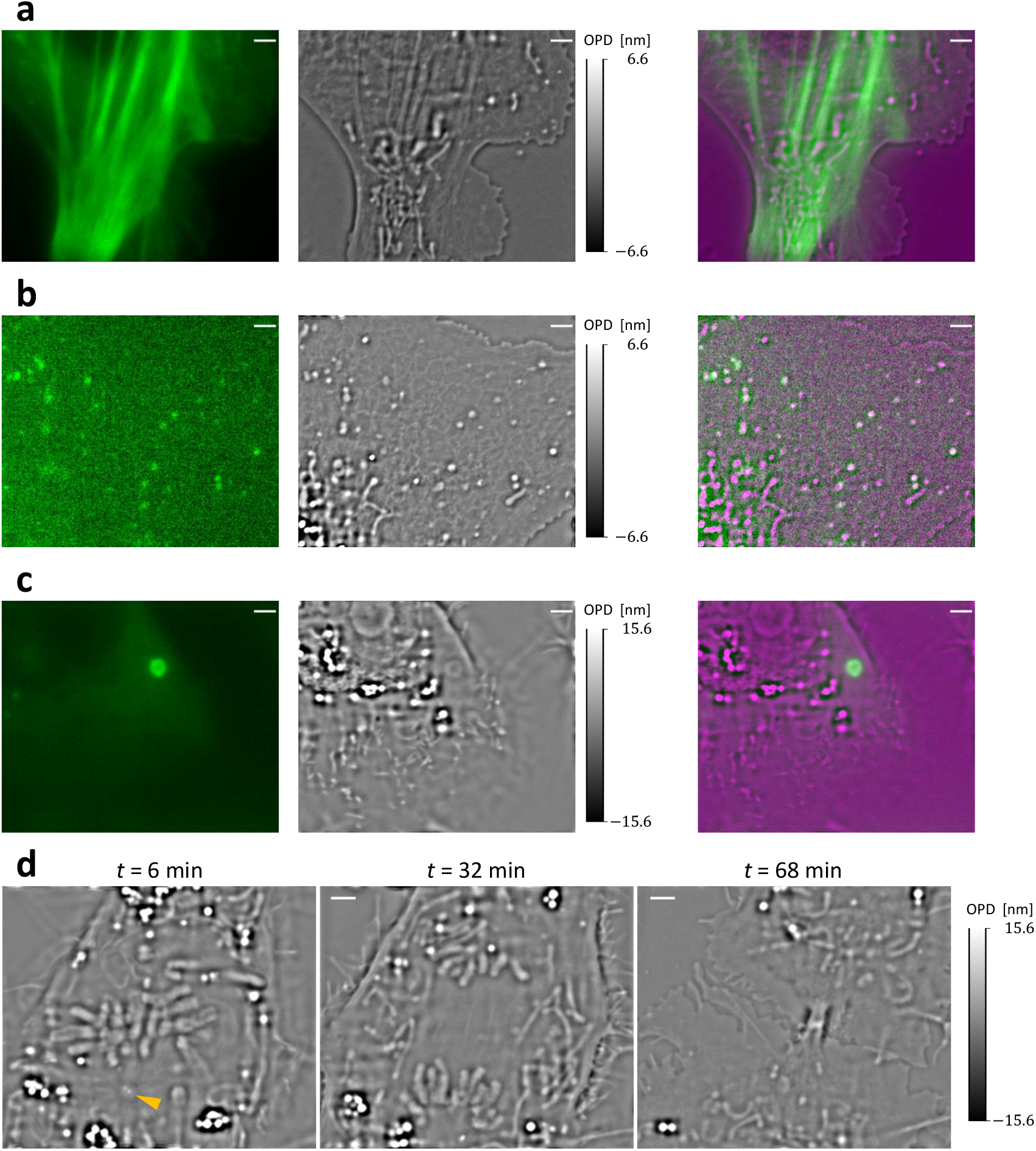
Observation of various organelles with the UpDPC system. **a-c**, Fluorescence image (left), UpDPC phase image (center), and composite image with phase using red color (right) of live cells stained with fluorescent proteins. **a**, PtK2 cell expressing mSG-βActin to visualize actin bundles. **b**, COS-7 cell expressing mSG-PTS1 to visualize peroxisomes. **c**, MDCK cell expressing G3BP1-mCherry to visualize stress granules. **d**, Time series of the retrieved phase shift of a COS-7 cell during mitosis. As shown in the left image, our system visualizes the two centrioles pointed by the yellow arrow during the M phase. Spindle fibers and chromosomes are also seen from prophase to anaphase, as shown in the middle image. After mitosis, the midbody structure between two daughter cells is captured, as revealed in the right image. The whole observation with a 30-ms time interval is shown in Supplementary Video 6. The scale bar shows 2 um.

Furthermore, due to UpDPC’s low phototoxicity, it is possible to examine processes that are highly sensitive to light-induced damage, such as cell division. Reactive oxygen species generated by fluorescence imaging can activate the spindle assembly checkpoint (SAC), halting cell division^26,27^. Therefore, fluorescence imaging conditions of cell division must be carefully considered, often requiring longer intervals between images, which decreases the temporal resolution, or reduced excitation light intensity, which increases noise. However, with UpDPC, we observed the entire process of chromosomal condensation, alignment, separation, and cytokinesis, leaving the midbody during over an hour of continuous imaging at 30 fps (Fig. 6d, Supplementary Video 5). Throughout this process, individual chromosomes, as well as centrosomes and surrounding spindle fibers, were visualized.

## Discussion

We introduced and implemented a high-speed, high-resolution, and high-sensitivity CPM technique, UpDPC nanoscopy, that can be easily extended from conventional microscopes. The required additional components are simple to integrate, with a total cost below 5,000 USD. Notably, UpDPC does not interfere with the epi-illumination path of inverted microscopes, allowing straightforward integration into complex epi-illumination configurations, such as structured illumination microscopy and optical tweezers. Simultaneous observation with fluorescence imaging is also achievable by effectively utilizing the near-infrared wavelength range (demonstrated in Fig. 6a-c). Additionally, this ease of implementation allows flexibility in adjusting setups for specific experimental needs, such as balancing resolution and photodamage. In this study, we prioritized resolution to achieve sub-100 nm rectangular gap separation and demonstrated high-speed long-term imaging with long-wavelength LEDs. UpDPC can be integrated into microscopes with stage-top incubators to capture the morphological changes of iPS cells differentiating into neurons, which are highly sensitive to phototoxicity (Extended Data Fig. 12). This convenient configuration, which extends commercial microscopes and whose optical system remains unchanged during imaging, parallels the successful adoption of techniques such as ZPC and DIC microscopy. Therefore, UpDPC is expected to be widely adopted in biological sciences.

We evaluated the performance of UpDPC in comparison with other QPI techniques, summarized in Extended Data Table 1. UpDPC demonstrated the ability to transmit spatial frequency information up to 2NA_obj_/*λ*, achieving resolutions twice the coherent diffraction limit. Among QPI techniques, it ranks among the highest spatial resolutions, surpassed only by Fourier ptychography, which requires many images. UpDPC can operate at the camera’s maximum frame rate and avoid motion artifacts caused by phase retrieval. Thus, UpDPC strikes an effective balance between spatial and temporal resolution, though this comes at the expense of FOV due to the need to retrieve phase from 2×2-pixel blocks. However, this sacrifice is reasonable given the role of new microscopy in expanding the horizon of observable phenomena. Future integration with gigapixel imaging techniques^28,29^ could further enhance UpDPC’s potential. Additionally, the phase retrieval in UpDPC is computationally efficient, allowing for real-time phase imaging on standard PCs through direct computation without the need for iterative methods. Furthermore, UpDPC does not require spatial coherence in the illumination source, eliminating speckle noise and enabling more robust imaging.

Label-free, high-resolution spatiotemporal imaging can offer new insights into cell biology. In this study, we successfully observed stress granules associated with G3BP1, recently reported to be invisible in QPI due to their refractive index similarity to the cytoplasm^30^. UpDPC revealed an internal sparse structure surrounded by a dense surface (Fig. 6c). This demonstrates that even structures with minimal refractive index contrast, on average, can still reveal density variations when observed at high resolution and high sensitivity. Furthermore, a modified QPI based on spatial light interference microscopy (SLIM)^31^ was used to visualize the dynamics of mitochondria, which are highly sensitive to phototoxicity by strong illumination for fluorescent imaging^32^. Similarly, high-resolution, label-free observation is expected to advance the understanding of photosensitive cell types such as iPS cells and early-stage embryos.

In modern cell biology, where the interplay among cellular components is increasingly emphasized, high-resolution imaging of entire cellular components is crucial. While fluorescence imaging offers the advantage of specific labeling of target structures, it can overlook interactions with the surrounding environment. Advanced techniques have been developed to visualize organelle contacts through multicolor fluorescence imaging^33^, but the drawbacks of fluorescence increase with the number of colors. In contrast, label-free UpDPC provides comprehensive information about the entire cellular environment. With recent advances in machine learning, the highly reproducible phase shifts of organelles could allow each component to be labeled based on unique patterns. The high throughput of UpDPC further facilitates the collection of dynamic data, enabling not only the analysis of spatial refractive index patterns but also their temporal evolution, opening new avenues for research in cell biology.

## Acknowledgments

We thank our lab members for their technical support and discussion, especially Mr. Rintaro Shimojo for contributing to the initial phases of this study and Mr. Lee Andre for English editing. We also thank Dr. Takuro Ideguchi, Dr. Kazumasa Takeuchi, Dr. Zenas C. Chao, Dr. Uri Manor, Mr. Katsuyuki Abe, Mr. Hiroshi Ishiwata, and Mr. Shinichi Hayashi for valuable advice and discussion. We would like to thank Ms. Junko Asada for technical assistance, Ms. Tomoko Furuya, and Ms. Manaho Kakiuchi for secretarial assistance. Y.I. is supported by the World-leading INovativeGraduate Study Program (WINGS) for the Forefront Physics and Mathematics Program to Drive Transformation (FoPM) of The University of Tokyo and the Japan Society for the Promotion of Science (JSPS) Doctoral Course (DC) Research Fellowship. This work is also supported by JSPS through KAKENHI grants (19H03394, 19H05794, 19H05795, 22H02798, 22H04926, 23K24060 and 24H02758 to Y.O.; 23KJ0643 to Y.I.), by the Japan Science and Technology Agency (JST) CREST grants (JPMJCR20E2, JPMJCR1852, JPMJCR24T2 to Y.O.) and the Moonshot R&D grant (JPMJMS2025-14) to Y.O.

## Author contributions

Y.I. and Y.O. conceived the project. Y.I. designed the phase retrieval scheme, conducted experiments, and analyzed the data. Y.I., M.K., and Y.O. designed the microscope. K.Y. fabricated the resolution evaluation targets. Y.I. wrote the manuscript draft. All authors edited the manuscript.

## Methods

### Microscopy setup

UpDPC was implemented on an inverted microscope (IX-83, Evident). The condenser lens setup included a universal condenser unit (U-UCD8 with IX-ADUCD, Evident) and a 1.4 NA oil-immersion top lens (U-TLO, Evident). A hyper red (peak wavelength of 660 nm; GH CSSRML.24, ams OSRAM), red (dominant wavelength of 625 nm; OSR5XNE3C1S, OptoSupply), and deep blue (peak wavelength of 445 nm; GD CSSRML.14, ams OSRAM) power LED were used for the transmitted illumination light source. This LED was equipped with an aluminum plate that we cut with an NC milling machine and then attached to the condenser lens using a 3D-printed holder. A quadrant polarizing filter was made from the linear polarizer film (extinction ratio of 9000:1 and polarization efficiency > 99.98%; XP42-18, Edmund optics). This film, fixed on an overhead projector (OHP) sheet, was placed in front of the condenser filter wheel. To prevent high-power unpolarized light from entering through the gaps in the polarizing film and to improve tolerances for film manufacturing, we printed a pattern of a cross and a disk on the OHP sheet twice using a printer (IM C3000, RICOH) set to maximum density. Since the OHP sheets also shift polarization, the polarizing film was positioned at the optical element closest to the sample. A 1.4 NA oil immersion objective lens (UPLSAPO100XO, Evident) was used as the objective lens. The polarization camera (CS505MUP, Thorlabs) was mounted with a magnifier made to compensate for the coarse-graining effect of the polarizing grids. The magnifier was constructed with a 30 mm cage system (Thorlabs) and two achromatic doublet lenses with focal lengths of 30 mm and 100 mm (AC254-30-A-ML and AC254-100-A-ML, Thorlabs). As a result, the effective pixel size is 20.7 nm, providing 3.8–5.7× oversampling relative to the Nyquist limit of the diffraction-limited resolution.

Except for Fig. 6, fluorescence images were acquired on the same IX83 microscope using a broadband LED light source (XT720S, Excelitas Technologies) and dichroic mirrors (U-FBNA for StayGold, U-FGNA for tetramethylrhodamine and mCherry, Evident). For Fig. 6, to simultaneously acquire fluorescence and quantitative phase images, a notch dichroic mirror (ZT656dcrb, Chroma), which reflects only the hyper-red LED for transmitted light, was mounted on the upper deck of the IX83 microscope. This was achieved using a mirror unit (91044 Laser TIRF for Olympus BX3/IX3 models, for 25mm filters, Chroma) inserted into the right side port (IX3-RSPC) and connected to the optical path leading to the polarization camera.

To reduce the intensity of the hyper-red LED light passing through the dichroic mirror, a bandpass filter (661 nm CWL, 25 mm Dia, 20 nm Bandwidth, OD 6 Fluorescence Filter, Edmund) was placed in front of the LED’s collimating lens. Fluorescence light passing through the fluorescence dichroic mirror in the filter wheel on the lower deck of the IX83 microscope was captured by a CMOS camera (ORCA-Fusion, C14440-20UP, Hamamatsu Photonics).

When aligning the fluorescence and phase contrast positions, a characteristic bright-field image was captured using the original LED light source of the IX83 microscope. Approximately 10 coordinates of distinctive structures were manually identified, and a 2×2 transformation matrix minimizing the sum of squared distances was calculated for correction.

The source intensity observation on the back focal plane of the condenser lens was observed by constructing the optical system shown in Extended Data Fig. 1c using a 50 mm focal length achromatic doublet lens (AC254-50-A-ML, Thorlabs) instead of the magnifying system.

The ZPC setup was implemented on the same IX83 microscope by combining a phase-contrast objective lens from the same series (UPLSAPO100XOPH, Evident) with a corresponding phase ring (U-PH3-S, Evident) and a dry top lens (U-TLD, Evident) on the same condenser unit.

The DIC setup was implemented using the same objective lens as in the UpDPC system along with the objective Nomarski prism (U-DICT, Evident) and the analyzer (U-AUT, Evident), in addition to the corresponding condenser Nomarski prism (U-DIC100, Evident) and the dry top lens on the same condenser unit. The polarizer attached to the condenser unit was aligned perpendicularly to the analyzer by fixing at the angle where the intensity value at a patternless area on a Siemens star glass was minimized. The bias phase retardation of the objective prism was adjusted to π/4 by rotating the bias retardation screw through one entire cycle and setting it at the angle where the intensity value was precisely halfway between the maximum and minimum.

For iPS cell observation, our UpDPC was implemented on an inverted microscope (ECLIPSE Ti2-E, Nikon) with a stage-top incubator and warming box system (Tokai Hit). A 0.52 NA dry condenser lens unit (Ti-C-LWD) and the same LED as above were used. The same polarizer film was sandwiched between two glasses (30mm Diameter Uncoated B270 Window, Edmund optics; 30mm Dia. 3mm Thick Uncoated, 1λ Fused Silica Window, Edmund optics) mounted with handmade 3D-printed parts insertable into the condenser turret. A 0.5 NA dry objective lens (Super Fluor 10X, Nikon) was used. The polarization camera, same as above, was mounted with a magnifier made with a 30-mm cage system (Thorlabs), objective lens (LUCPLFLN20X, Evident), and achromatic doublet lens with a focal length of 100 mm used above.

All microscopes were controlled by custom scripts written in Python. The microscopes were connected to the Micromanager^34,35^ using the appropriate driver, and the Micromanager was controlled from Python using the Pycro-Manager module^36^. The Thorlabs camera was controlled directly from Python using the Thorlabs camera SDK to maximize image acquisition speed. The code is uploaded to https://github.com/inutsuka-yugo/UpDPC.

### Protocol of microscopy setup and observation

When placing samples, we put a drop of immersion oil (IMMOIL-F30CC, Evident) on the objective lens while caring not to make air bubbles, set the sample on it, and carefully put the immersion oil on the sample enough to cover the condenser lens on it. The objective lens should be slowly moved closer to the sample bottom in order not to create bubbles. After that, we move the condenser lens closer to the sample top until the condenser lens is covered with the oil by confirming the increase of the reflected light from the oil. We check the pupil plane by setting the optical path to eyepiece mode and seeing the sample without the eyepiece.

For positioning the lenses to the Köhler illumination, we close the aperture stop so that our eyes can see the sample easily and focus the sample through the eyepiece utilizing the XY stage. After finding the focus, we fully open the aperture and close the field aperture to locate the optical axis. This optical axis is matched to the center of the FOV. At the same time, we carefully move the condenser lens up and down and fix the position where the polygonal vertices of the field diaphragm are visible. Then, we can put the additional power LED. We darken the room after this setup to remove the noise.

The camera’s rotation angle is corrected to the same as the XY stage. We use motionless objects, such as debris stuck to the glass. We bring this object to the edge of the FOV and rotate the camera so that the XY stage motion is parallel to the edge.

Systematic noise information is acquired by capturing an image series (100 frames in this case) at odd-numbered FOVs (usually 5 FOVs), where almost all areas consist of a glass surface without additional phase objects. At the same time, the exposure time is adjusted to nearly fulfill the camera’s dynamic range. After that, a temporal average excluding outliers (defined here as deviations greater than two standard deviations from the median) is calculated for each pixel in the image series of each FOV. Subsequently, the median of the temporally averaged images at odd-numbered FOVs is computed. This strategy effectively removes the slight additional objects on the glass surface. For example, the 10-pixel diameter debris can be ignored by taking the median of 11-pixel right and 11-pixel upward. As long as the majority of frames in each pixel correspond to the glass surface, a clean background image, denoted as *I*_0_(***x***) later, can be successfully created.

### Illumination source function determination

To determine the illumination intensity pattern at the back focal plane of the condenser during phase retrieval, a Bertrand lens was added instead of a magnifying lens to observe the illumination pupil plane through a uniform sample. Four sample conditions were used: [1] a glass slide alone, [2] L15 medium sandwiched between a glass slide and a coverslip with a thickness of 30 μm, [3] L15 medium sandwiched between a glass-bottom dish and a coverslip with a thickness of approximately 1 mm, and [4] air sandwiched between a glass-bottom dish and a coverslip with a thickness of approximately 1 mm.

When fitting the illumination pattern, the polarized camera images were first normalized by dividing each polarized image by each average intensity to correct for the light path transmission rate in each polarization direction and then summed. To reduce the computational cost of the 2D fitting, these images were cropped to a smaller area where the entire illumination pattern was visible. The fitting function was based on a pattern excluding the cross and central circle from a 2D radially symmetric cosine function, blurred by the optical system’s point spread function (PSF). The Airy function of the PSF was approximated as a Gaussian, and the following function was used:

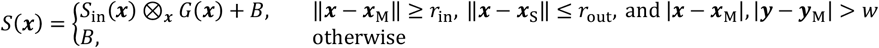

Where 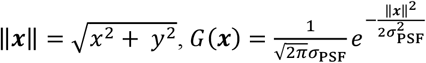, and 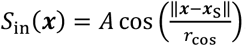. The cosine function in this *S*_in_ could be replaced by a quadratic or quartic function and a good fit was observed, but the cosine function consistently gave the best results with the number of parameters.

Initially, to perform a coarse fit, the image of the glass slide condition [1] was binned into an 8×8 grid, and the fitting conditions were evaluated using Python’s lmfit module. This function was fitted to the image 10 times with all fitting methods without providing the Jacobian, using visually estimated values as initial conditions. The random seeds were fixed at 0, 1, 2, …, 9 to ensure reproducibility in stochastic optimization methods. After comparing the methods based on the results (see Extended Data Fig. 3), we chose to use the ‘differential_evolution’ method. This method had the shortest computation time among the best comparable methods (‘dual_annealing’, ‘powell’, and ‘differential_evolution’), and it also resulted in the smallest mean value for the reduced chi-square.

Next, for all four sample conditions [1-4], a coarse fit was similarly performed on the binned images, and the resulting values (scaled by a factor of 8 for parameters of ***x***_M_, ***x***_S_, *r*_in_, *r*_out_, *w, r*_cos_, and *σ*_PSF_) were used as initial values for fine-tuning. Subsequently, the parameters that should be constant regardless of the sample (*r*_in_, *w*, and *r*_cos_) were fixed at the average values across the four conditions, and another fine-tuning was performed.

For these parameters, since the NA_ill_ should be 1.4 for the glass slide condition [1], approximately 1.33 for the 1 mm L15 medium condition [3], and approximately 1.0 for the air condition [4], with *r*_in_ = 0 corresponding to NA_ill_ = 0, a plot of *r*_out_ vs. NA_ill_ was created. The ratio between pixel and NA was determined using least-squares fitting, and the final values were set as *r*_in_ = 0.330, *w* = 0.073, and *r*_cos_ = 1.435 (in units of NA).

### Phase retrieval

The propagation model is described in Equations (1)-(5)in Supplementary Note 1. However, there are individual differences in each pixel and dust on the sensor in the camera, so the ideal obtained intensity is

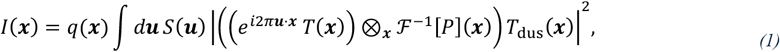

where *q*(***x***) is the quantum efficiency on each pixel and *T*_dus_(***x***) = *a*_dus_(***x***)exp(*iϕ*_dus_(***x***)) is the complex amplitude transmittance of the dust near the imaging sensor in the camera. We can easily see that the phase shift near the camera does not affect the intensity and

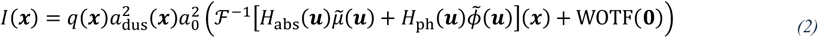

after the weak object approximation. Simultaneously, the obtained intensity *Î* includes the noise *ξ* related to the ideal intensity *I*, as *Î*(***x***) = *I*(***x***) + *ξ*(***x*;** *I*). This noise includes dark current, read noise, and shot noise. The most dominant part is the shot noise following a Poisson distribution related to the photon number corresponding to the intensity. Thus, this shot noise has a variance of a constant times *I*(***x***). In our setup, the photon number is sufficiently large, and we confirmed that the obtained intensity distribution for the static sample is a Gaussian distribution.

We can get the background image 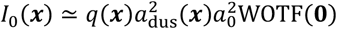 for a constant *μ* and *ϕ* as written in the previous section. The shot noise term can be small enough to ignore compared with the signal by averaging the long-time images. By dividing the obtained images by this background, we can acquire the corrected image 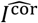 with the scaled noise: 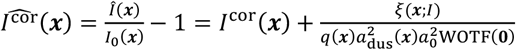 with a simple image transferred from the sample, 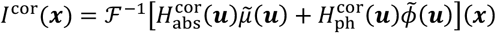, where 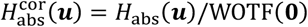 and 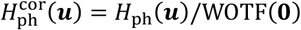.

Then, we estimate the most probable sample phase shift from the obtained image. In our setup, the image can be regarded as a mixture of images illuminated by four different light source patterns. We combine this information with Bayesian inference, finding a phase shift map Φ = {*ϕ*(***x***)}_***x***_ from the obtained images 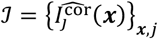 by maximizing the posterior distribution Pr(Φ|ℐ). Here and after, *j* denotes the intensity pattern index and *j* ∈ {1,2,3,4} in our setup. Using Bayes’ rule, this posterior distribution is calculated from the prior distribution and likelihood function as 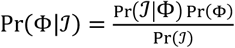,so we look for the phase shift map Φ maximizing the product of Pr(ℐ|Φ) Pr(Φ). Pr(ℐ|Φ) for each ***x*** is a Gaussian distribution with a variance of *ξ*(***x***;*I*) divided by 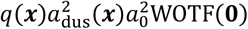, defined as *σ*_*I*_(***x***). To calculate the optimal *ϕ*(***x***) analytically, we assume Pr(Φ) is a Gaussian distribution with a variance of *σ*_*ϕ*_ for each ***x***. Then, 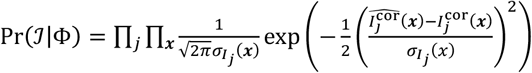 and 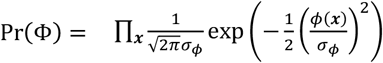, so we can consider the maximum of Pr(ℐ|Φ) Pr(Φ) with the logarithm as

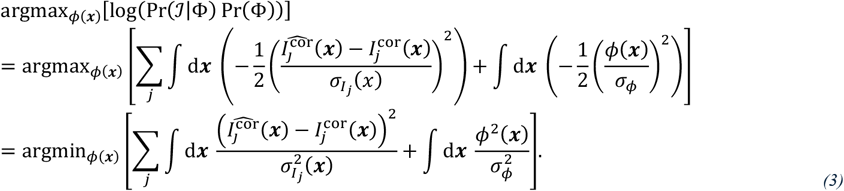

This argmax can be obtained as the solution where the functional derivative of the target function with respect to φ(***x****’*) is zero in the entire ***x****’*-space. This functional derivative is calculated as 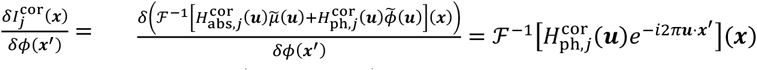, so this equation is 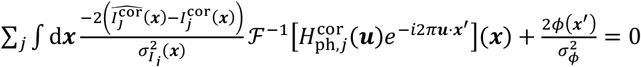. Taking Fourier transform from ***x****’*-space to ***v***-space and considering the minimum on ***v***-space simplifies this equation to 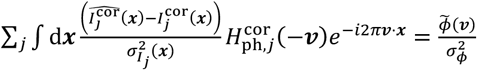 because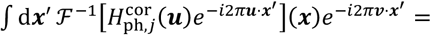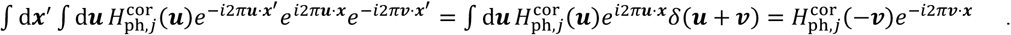.

Therefore, the equation is

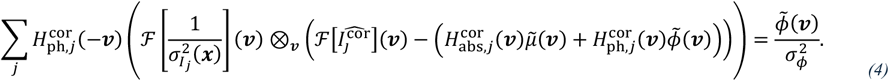

We can also derive the equation for absorption as

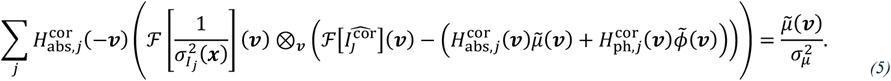

These equations can be analytically solvable by approximating *σ*_*Ij*_(***x***) to a constant *σ*_*I*_ and regarding 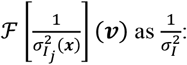

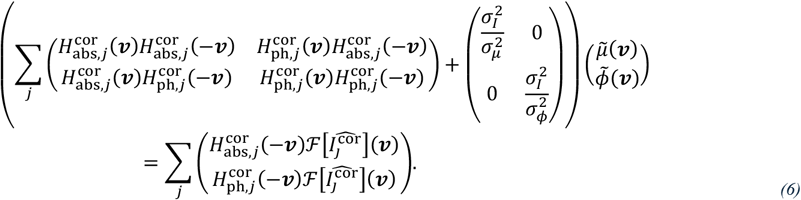

The 2×2 matrix on the left-hand side has a non-zero determinant with a non-zero *σ*_*I*_; thus, this equation has a solution. Also, 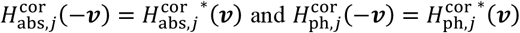, so this matrix becomes a positive Hermitian matrix 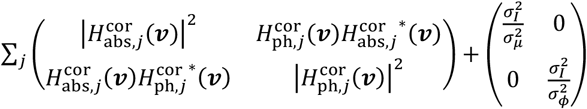 with positive diagonal components. We used this solution as the phase retrieval result.

This solution is identical to the Tikhonov regularization result when 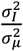 and 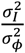 are regarded as the regularization parameters. These values are difficult to determine, so we tried values to the power of 10. If we set *σ*_*ϕ*_ too small, the noise on the image would be amplified, while if we chose too big *σ*_*ϕ*_, the original structures inside the cell would be erased. For optimal results, we used the 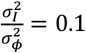 for cells and 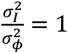 for the gliding assay, based on visual judgement. The value of <u>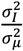</u> does not affect the retrieved phase very much, so we used the same value as 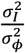. Thus, in real-time phase retrieval with observation, we ignored the absorption term and calculated the phase shift as 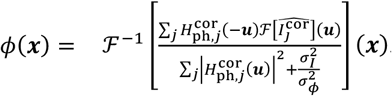.

Note that if the prior information about the phase shift probability distribution is known, using that distribution can improve this MAP estimation. However, we cannot analytically solve the optimum values in most probability distributions and need numerical optimization. It takes much more time than the direct computation method shown here to perform these computations, challenging applications to high-speed imaging data.

The phase retrieval code has been modified and implemented with reference to the data structure of the Waller-Lab DPC code^21,37^. The implementation is available at https://github.com/inutsuka-yugo/UpDPC.

### Cell culture and transfection

The COS-7 and MDCK cells were obtained from the Health Science Research Resources Bank in Osaka. The HeLa cells were a kind gift from Miho Ohsugi.^38^ PtK2 cells were obtained from the JCRB Cell Bank. These cells are maintained in DMEM (Gibco, Thermo Fisher Scientific) with 10% fetal bovine serum (Gibco, Thermo Fisher Scientific) at 37°C under 5% CO_2_. For the short-time observation of COS7 cells of Fig. 1e and Extended Data Fig. 2, cells transfected with TransFectin (Bio-Rad) were placed on glass coverslips (No. 1S, Matsunami) inside the plastic dish to realize the high illumination NA. Just before observation, we attached the glass slide (S2111, Matsunami) with 30-µm double-sided tape (No.5603, Nitto) and sealed it with silicone (DENT SILICONE-V, regular type, SHOFU). For simplified cell observation, cells reverse transfected with TransFectin were plated on a 50 mm diameter glass-bottom dish (P50G-1.5-14-F/H, MatTek), which does not interfere with the condenser lens. Before observation, the dish was sealed with the same silicone after placing the coverslip on top. When using the more common 35 mm diameter glass-bottom dish (P35G-1.5-14-C, MatTek), the plastic rim that could interfere with the condenser was removed using scissors. Before imaging, the medium was changed to Leibovitz’s L15 medium without phenol red (Gibco, Thermo Fisher Scientific) with 10% fetal bovine serum.

Human iPS cells (253G1 F12 strain) were obtained from RIKEN Cell Bank. A neurogenic transcription factor, neurogenin 2, was knocked into these cells’ safe harbor locus AAVS1 under the control of a tetracycline-inducible promoter. These cells were maintained in AK02N medium (StemFit, Takara Bio) at 37°C under 5% CO_2_. For 24 hours after each passage, ROCK inhibitor (Y-27632, Nacalai Tesque) was also added to the medium. On the day before the observation, iPS cells were plated on the glass-bottom dish (No. 1.5, MatTek) coated with iMatrix-511 (Clontech, Takara Bio) and cultured with induction medium composed of DMEM/F12 (Gibco, Thermo Fisher Scientific), N2 Supplement with Transferrin(Holo)(×100) (FUJIFILM Wako Pure Chemical Corporation), 1% MEM Non-Essential Amino Acids Solution (100X) (Gibco, Thermo Fisher Scientific), 1% GlutaMAX™ Supplement (Gibco, Thermo Fisher Scientific), and 2 ug/mL Doxycycline (Clontech, Takara Bio). ROCK inhibitor was added to this induction medium on this day. Before imaging, the medium was changed to the induction medium, whose doxycycline concentration is 5 ug/mL and without ROCK inhibitor.

For plasmids, er-(n2)oxStayGold(c4)^25^ (a gift from Atsushi Miyawaki) was used to label the endoplasmic reticulum. Additionally, monomeric StayGold (mStayGold, QC2-6 FIQ)^39^ (a gift from Atsushi Miyawaki) was used to label β-actin (referencing Addgene #21948), peroxisomes (peroxisomal targeting signal-1, PTS1), and Golgi apparatus (Giantin, Addgene #85048). For visualizing stress granules during arsenite stress, G3BP1-mScarlet was constructed by inserting human G3BP1 into mScarlet-i (Addgene #85044, a gift from Dorus Gadella) through infusion cloning.

### Nanofabrication and observation of handmade Siemens star target

As the reported method^40^, the handmade Siemens star target was fabricated with electron-beam lithography and dry etching on a fused-silica glass substrate (70 mm × 30 mm, VIOSIL-SQ, Shin-Etsu Chemical). The validation of the fabricated target was performed both laterally and axially.

Lateral validation was confirmed by scanning electron microscopy (SEM; Versa 3D Dual Beam, FEI). A vacuum was applied after depositing approximately 3 nm of Pt-Pd and loading the sample. The beam acceleration voltage was set to 5.0 kV, and the Z-axis was automatically corrected. As a result, patterns with a minimum width of approximately 50 nm were observed (Extended Data Fig. 5).

Axial validation was performed using white light interferometry (VK-X3050, Keyence). However, due to the difficulty in accurately measuring the small width of the Siemens star pattern, a reference pattern with a width of approximately 5 μm, fabricated under the same conditions (Extended Data Fig. 6, left), was created, and its depth was measured. The average depth was calculated by fitting the height probability density of the image data, spanning 283 μm × 212 μm within a 1024 × 768-pixel image, to a sum of two Gaussian functions, using the difference between the two means as the measured depth (Extended Data Fig. 6, right).

## Modulation transfer function (MTF) analysis

### 1. MTF derivation

In optical systems, to transfer images linearly or in fully incoherent light propagation, the resulting image *I*_res_ can be modeled as a convolution of the original image *I*_orig_ with the system’s point spread function (PSF): *I*_res_(***x***) = *I*_orig_(***x***) ⊗_***x***_ PSF(***x***). By performing the Fourier transform, this relationship can be simplified to the product as 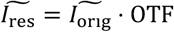, where the OTF is the Fourier transform of the PSF called the optical transfer function. If the original image is the arbitrary sinusoidal wave 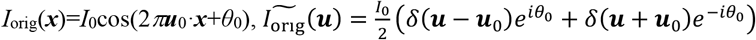, the resulting image is

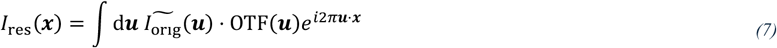

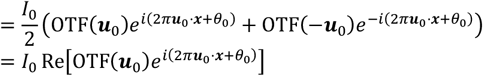

since the PSF is real and thus 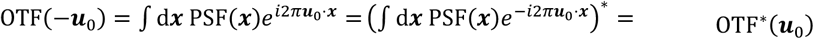. By separating the OTF with its amplitude and phase term as OTF = MTF*e*^*i*PTF^ with the real functions of the MTF and the PTF (phase transfer function), we can further calculate the resulting image as

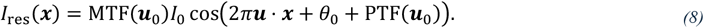

Therefore, the MTF indicates the contrast magnification for the given spatial pattern.

### 2. Pseudo-MTF analysis

We evaluated the spatial resolution by observing the phase shift of the periodic pattern on the circumference of the Siemens star with the UpDPC system, as is frequent practice in QPI papers. We assumed the concave-convex pattern is blurred in the nanoscope, as in many other microscopes. In other words, let *l*_*r*_ be the coordinate on the circle of radius *r* in the Siemens star, and let *ϕ*_orig_(*l*_*r*_) be the actual phase shift profile on this *l*_*r*_. We assumed the phase shift profile *ϕ*_res_(*l*_*r*_) is obtained by convolution with

PSF_*ϕ*_(*l*_*r*_) as *ϕ*_res_(*l*_*r*_) = *ϕ*_orig_(*l*_*r*_) ⊗_*l*_ PSF_*ϕ*_(*l*_*r*_). The spatial resolution is then evaluated quantitatively by calculating 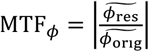. Here, *ϕ*_orig_(*l*_*r*_) can be well described by Fourier series expansion as 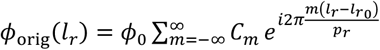 with the coefficient 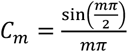, thanks to the periodicity, where *ϕ*_0_ is the spoke phase shift, 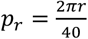 is the period of the fringes on a circle with radius *r*, and 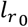 corresponds to the center of some gap on 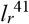 . Thus, 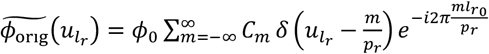 indicating that 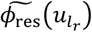 has information at spatial frequencies of integer multiples of 1/*p*_*r*_. However, as fabricated by dry etching, the circumferential cross-section of the Siemens star does not have a perfect square wave profile, with slightly rounded corners. Accordingly, we focused on the component with the largest contribution at the frequency 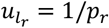 for each circumference and obtained the value of MTF_*ϕ*_. That is, the value of 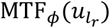 was determined to be

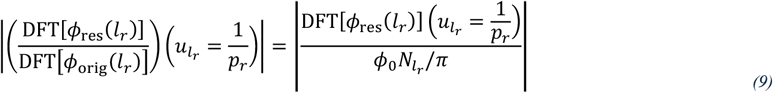

at the radius *r* where 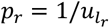. Note that DFT is the discrete Fourier transform, and 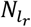 is the number of data points for *ϕ*_res_(*l*_*r*_) corresponding to the delta function in the DFT. This value can be understood intuitively as the ratio of how much of the wave component closest to the concavity pattern of each phase profile was transmitted. Similarly, for ZPC and DIC images, we calculated the pseudo-MTF (pMTF) value as

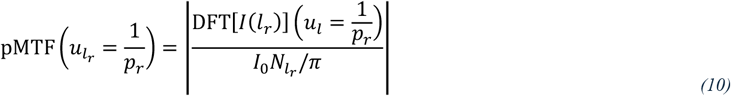

for the 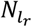 data points of intensity values *I*(*l*_*r*_) on a circumference of radius *r*, where *I*_0_ = 1.

### 3. Microscope observations of Siemens-star target

The Siemens star pattern was pressed down with a coverslip, either not putting anything or after adding MilliQ water. The boundary of the coverslips was sealed with silicone. During observation, five samples that were free of dust and had no visible pattern defects were selected for each condition. For each field of view, 36 images were captured at 0.1 μm intervals, within a range of ±5 frames relative to the visually focused position. The exposure time was adjusted so that the maximum brightness value in the field of view was around 3900, just below the maximum value of 4096 for a 12-bit camera. As a result, exposure times of 110 ms for ZPC, 55 ms for DIC, and 8 ms for UpDPC were used.

After imaging, for each Z-plane’s 36 images, the mean and standard deviation were calculated for each pixel. Any pixel whose absolute deviation from the mean was more than three times the standard deviation was considered an outlier and excluded. An average image was then generated. Next, the focal plane was selected for each field of view by taking the Z-position where the log|DFT[image]| averaged over each image had the maximum value. Subsequently, the center position of the Siemens star in each field of view was determined by image processing and manual fine-tuning. For the ZPC and UpDPC images, a Gaussian blur with a standard deviation of *σ*_blur_ was applied, and pixels with a deviation from the mean (or its negative for UpDPC) greater than *n*_std_ times the standard deviation were extracted, isolating the Siemens star partially with the outermost region being centrally symmetric. Then, binary images were dilated by *k*_dilate_ to create a centrally symmetric and internally uniform region, and the centroid of this region was set as the star’s center. For the DIC images, similarly, a Gaussian blur with a standard deviation of *σ*_blur_ was applied, and pixels with an absolute deviation from the mean larger than *n*_str_ times the standard deviation were extracted, isolating regions with high brightness perpendicular to the prism shear direction. Edges were detected using a Canny detector, and Hough line transform was used to generate lines, most of which passed through the center. Finally, the intersections of these lines were calculated, and the centroid of these intersections was defined as the star’s center. The parameters *σ*_blur_, *n*_str_, and *k*_dilate_ were chosen based on visual inspection to produce the best results. The code for these procedures has been uploaded to https://github.com/inutsuka-yugo/UpDPC.

As discussed in the previous section, the estimation of the phase difference in the Siemens star was based on the assumption that the Siemens star pattern can be expressed as 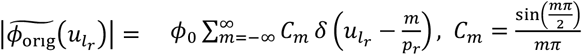. By directly assuming that *ϕ*_res_ = *ϕ*_orig_, the phase difference was estimated at each radius *r*, where 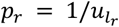, using 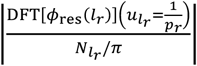. The design values for comparison were defined as |2π (*n*_medium_ – *n*_glass_) × *d*_WLI_ / *λ*|, where *n*_medium_ is the refractive index of the surrounding medium (1 for air and 1.33 for MilliQ water), *n*_glass_ = 1.45 is the refractive index of the glass used for the Siemens star, *d*_WLI_ is the average height measured by white light interferometry (WLI), and *λ* = 445 nm is the central wavelength of the illumination. The code for these calculations has also been uploaded to https://github.com/inutsuka-yugo/UpDPC.

### Gliding assay

In the gliding assay, microtubules, a small amount labeled with tetramethylrhodamine (42.6 nM in non-label 180 nM), are run on kinesins K560 (mouse) attached to the glass with the His tag. The samples were prepared by adding His-tag antibody (1/10), 0.9 uM kinesin with His tag on C-terminal, 1 mg/mL casein in PEM80, assay buffer (PEM80, 1mM ATP, 0.2 mg/mL casein, 10 *μ*M taxol, 1.0% 2-mercaptoethanol, 0.2 mg/mL glucose oxidase, 40 *μ*g/mL catalase, 1 mM glucose, final pH ∼ 6.9), and microtubules in this order to a channel consisting of a glass slide and cover glass sandwiched between 30 μm double-sided tape and sealing with commercial nail polish.

### Silica beads observation

To determine localization accuracy, fluorescent silica beads with a diameter of 500 nm and a plain surface (sicastar®-greenF, micromod Partikeltechnologie) were observed in a glass chamber similar to that used for the gliding assay, sealed in silicone oil with an adjusted refractive index.

The refractive index of the silicone oil was adjusted by mixing two commercially available products (KF-50 and KF-53, Shin-Etsu Silicone). The RIs of the silicone oils were adjusted by mixing two commercial products (KF-50 and KF-53, Shin-Etsu Silicone). The RIs of the two silicone oils were measured with a pocket refractometer (PAR-RI, Atago) and found to be 1.428 and 1.485 at 25°C. We also confirmed that the RIs of their mixtures correlated linearly with the mixing ratio. By measuring the phase shift of silica beads in this silicone oil, the RI of silica beads was confirmed to be ∼1.44.

We used silicone oil with RI of 1.448 mixed with 636.8 mg of KF-50 and 388.6 mg of KF-53 in our experiments to evaluate the localization accuracy under the condition of 500 nm × (1.448 − 1.44) ∼ 4 nm optical path length difference.

### Vesicle tracking and step detection

In vesicle tracking, the center position of each vesicle is determined by Gaussian 2D fitting. Then, the nearest vesicles are connected using the k-d tree structure. These (position, frame) data are trimmed to 500-ms sub-trajectories to fit trajectories to the line and decomposed parallel and perpendicular components. To search for one-directional and stepwise trajectories, we filtered the trajectory with some conditions, such as maximum distance above 100 nm, maximum forward velocity above 2 nm/ms, maximum backward velocity less than 1 nm/ms, and velocities in 60% frames less than 1.25 nm/s. Then, the trajectories are analyzed with the fine-tuned step-finding algorithm for low signal-to-noise ratio observation^42^.

